# The replication of the human respiratory syncytial virus in a T cell line has multiple ineffective steps

**DOI:** 10.1101/2020.12.28.424605

**Authors:** Ricardo de Souza Cardoso, Ana Carolina Lunardello Coelho, Bruna Laís Santos de Jesus, Brenda Cristina Vitti, Juliano de Paula Souza, Rosa Maria Mendes Viana, Marjorie C. Pontelli, Tomoyuki Murakami, Armando Moraes Ventura, Akira Ono, Eurico Arruda

**Author notes:** Contributed equally to this study (Akira Ono, Eurico Arruda).

## Abstract

Human respiratory syncytial virus is the most frequent cause of severe respiratory disease in children. The main targets of HRSV infection are epithelial cells of the respiratory tract and the great majority of the studies regarding HRSV infection are done in respiratory cells. Recently, the interest on respiratory virus infection of lymphoid cells has been growing, but details of the interaction of HRSV with lymphoid cells remain unknown. Therefore, this study was done to assess the relationship of HRSV with A3.01 cells, a CD4^+^ T cell line. We found by flow cytometry and fluorescent focus assay that A3.01 cells are susceptible but virtually not permissive to HRSV infection. De-quenching experiments revealed that the fusion process of HRSV in A3.01 cells is reduced in comparison to HEp-2 cells, an epithelial cell lineage. Quantification of viral RNA by qPCR determined that the replication of HRSV in A3.01 cells was modest. Western blot and quantitative flow cytometry analyses demonstrated that the production of HRSV proteins in A3.01 was significantly lower than in HEp-2 cells. Additionally, we found by fluorescence in situ hybridization that the inclusion body-associated granules (IBAG’s) are almost absent in HRSV inclusion bodies in A3.01 cells. We also assessed the intracellular trafficking of HRSV proteins and found that HRSV proteins co-localized partially with the secretory pathway in A3.01 cells, but these HRSV proteins and viral filaments are present only scarcely at the plasma membrane. HRSV infection of A3.01 CD4^+^ T cells is virtually unproductive as compared to HEp-2 cells, with virion production hampered by low fusion, hypofunctional inclusion bodies, altered trafficking of viral proteins to the plasma membrane.

## Introduction

Human respiratory syncytial virus (HRSV), of the family *Pneumoviridae*, is a common respiratory pathogen that circulates worldwide and a major cause of serious lower respiratory tract disease, mainly bronchiolitis in children, but also severe disease in the elderly [1]. HRSV infects mainly epithelial cells of the respiratory tract [1], but has also been detected in non-respiratory tissues and cells [2][3]. In that regard, HRSV and other respiratory viruses have been detected in tonsillar tissues and respiratory secretions from children with tonsillar hypertrophy without symptoms of acute respiratory infection [4], suggesting that HRSV may infect secondary lymphoid tissues. In addition, HRSV antigen has been detected in circulating T lymphocytes, and thus it is conceivable that it may affect immune function [5].

The HRSV genome is a single-stranded RNA with 10 genes that encode 11 proteins [1]. Virus entry is mediated by the G protein that binds to the host cell [6], resulting in the fusion between viral envelope and cell membrane, mediated by the HRSV F protein [7,8]. It is also known that HRSV enters cells by macropinocytosis, a process in which nucleolin participates as virus receptor [9][10]. HRSV replication involves production of inclusion bodies [11], where viral proteins L, P, N, M, and M1,2 are present [12–14]. However, the subsequent HRSV assembly process is not entirely understood. It has been shown that the viral glycoproteins follow the secretory pathway [15–18] to reach the plasma membrane but also utilize Apical Recycling Endosome (ARE) machinery during this process [19]. In contrast, the trafficking of HRSV non-glycosylated proteins during virus assembly has remained quite unclear. It is known that HRSV M protein forms dimers prior to participating in the formation of viral progeny [20], and that this protein has affinity to endomembrane system [21]. HRSV N protein coats the virus genome and forms inclusion bodies, an essential role in which it is helped by the P protein [22]. The inclusion bodies (IB’s) of the HRSV are places where the viral replication and transcription occur; in addition, the IB’s are responsible for stabilizing the viral mRNAs that confers an efficient protein translation process [14]. HRSV N protein was also observed to co-localize partially with the Golgi [23]. Consistent with these findings, recently we have shown that in HEp-2 cells the M and N engage partially with secretory pathway and with the retromer complex [24]. Furthermore, HRSV P protein was found partially co-localizing with endosomal vesicles [25]. Together, these findings suggest the possibility that the non-glycosylated proteins of the HRSV reach the assembly sites at least partially using secretory and/or endosomal pathways. The steps of viral assembly and budding take place at the plasma membrane of the infected cells [21,26,27]. It is known that HRSV F protein cytoplasmic tail is pivotal in the virus budding process [7,28]. Finally, HRSV budding process results in a filamentous viral particle, an event dependent on Rab11-FIP2 protein but not Vps4 [29].

Knowledge on HRSV-cell interactions has been accumulating through studies that use models based on epithelial cell cultures susceptible and permissive to HRSV infection. Nonetheless, virus-host cell interactions may be different between cells of epithelial origin like HEp-2, in which virus progeny production results in cell death, and lymphomononuclear cells, in which HRSV may cause long-term or persistent infection. Previous studies have shown that HRSV infects CD4^+^T cells [5] and Breg cells of neonates [30], which may enhance HRSV disease. Very little is known about HRSV replication and progeny production in lymphoid cells, and hence the present study was done to investigate intracellular HRSV assembly and replication in A3.01 cells, which belong to the lymphoid cell lineage.

## Material and Methods

### Cells and viruses

A3.01 cells, obtained from the AIDS Research Reagent Program, were maintained in Rosswell Park Memorial Institute (RPMI) culture medium, supplemented with 10% of fetal bovine serum (FBS) and 1% of antimycotic/antibiotic. HEp-2 and Vero cells were acquired from ATCC and maintained in MEM with 10% of BFS in 1% of antimycotic/antibiotic. The virus used for this study was HRSV A long strain (ATCC), propagated in HEp-2 cells and titrated in Vero cells, following routine agar-based plaque assays. The multiplicity of infection (MOI) used for the infections are specified in the experiment results.

### Antibodies

The antibodies used were FITC-conjugated mouse monoclonal anti-RSV N (MAB 858 3F, Millipore), mouse monoclonal anti-RSV F (MAB8262X, Millipore), mouse anti-CD63 (Clone H5C6 RUO, BD Pharmingen), mouse polyclonal anti-RSV M [12], rabbit polyclonal antibodies anti-TGN46 (ABT95, Millipore), anti-Giantin (PRB-114C-200, Covance), anti-SNX2 [31], anti-Lamp-1 D2D11 XPR (9091T, Cell Signaling); mouse monoclonal anti-EEA1 (610457-BD), goat anti-RSV (Abcam ab20745); mouse monoclonal anti-β-actin (sc 47778), Alexa Fluor 594-conjugated goat anti-rabbit (ab150080, Abcam), Alexa Fluor 647-conjugated goat anti-mouse (ab150115, Abcam), HRP-conjugated goat anti-rabbit (656120 Invitrogen), HRP-conjugated rabbit anti-goat (305 035 003, Jackson ImmunoResearch), and HRP-conjugated rabbit anti-mouse (A9044, Sigma).

### Flow Cytometry

HEp-2 or A3.01 cells were infected with HRSV, and the harvest occurred 24, 48, and 72 hours post infection, when cells were fixed with 4% PFA for 20 minutes. After that, cells were permeabilized and incubated with FITC-labeled mouse anti-RSV Nucleoprotein (Millipore), followed by washes with PBS containing 1% BSA. The cells were analyzed in a BD LSR Fortessa flow cytometer; the results were analyzed in the Flow Jo software. The gating strategy used to perform the single cell analysis for N-positive population to establish the mean fluorescence intensity was the 2D plot, in which the plans were fluorescence signal for N versus forward scatter. Experiments were done three times independently.

### Biobond and Poly-Lysine coverslips treatment

Coverslips were treated with 0.1 N HCl in 100% ethanol, then in 100% alcohol, air-dried, and then treated with 4% Biobond (Koch Electron microscopy) in acetone for 4 minutes, followed by a wash in distilled water and air-drying. The same procedure was used for treatment with Poly-Lysine, except for 20 µL of poly-lysine was used for 1 hour and then coverslips were air-dried. T cells were deposited onto pre-treated coverslips and incubated stationary for 3 hours, followed by fixation in 4% PFA for 20 minutes, washing, in PBB 1X and testing by immunofluorescence or fluorescent in situ hybridization (FISH).

### Immunofluorescence

After fixation, A3.01 or HEp-2 cells on coverslips were permeabilized in 0.01% Triton X-100 in PBS for 15 minutes, and washed five times in 1X PBS. The cell preparations were incubated with the appropriate primary antibody for 1 h in a humidified chamber at 37°C. After that, preparations were washed five times in PBS, incubated with the appropriate secondary antibody along with DAPI (Sigma) for 1 h, and finally washed in PBS. The coverslips were mounted on slides with Flouromount™ and analyzed by confocal microscopy in a Leica SP5 or a Zeiss 780 microscope. To perform analysis of quantity and size of the inclusion bodies in A3.01 and HEp-2 cells, the immunofluorescence images were subjected to Image J software analysis. Using the tool Analyze>Analyze Particles, it was possible to determine the area and number of the N-positive structures. The same procedure was done for measuring the size of the A3.01 cells, but in this case, after the analysis, the mean of the area of the A3.01 cells was established by conventional mathematic method.

### Fluorescent in situ hybridization

HEp-2 or A3.01 cells were seeded on coverslips treated or not with Poly-Lysine, and 24 and 48 hours post-infection were fixed with 4% PFA and tested by FISH as per published protocol [14], slightly modified. Briefly, fixed cells were permeabilized with 0.1% Triton X-100 in 1% BSA-PBS and then treated with free streptavidin for 1 h, re-fixed in 4% PFA, washed in PBS, incubated with the hybridization mix (1µM biotinylated poly T probe, 35% formamide, 5% dextran sulfate, herring sperm DNA in 2 X SSC2) at 37°C for 3 h, followed by serial washes in 2 X SSC, SSC, and PBS. After that, coverslips were incubated with Alexa Fluor 594 streptavidin, and then with FITC-conjugated mouse anti-RSV N (Millipore) for 1 h. Finally, the cells were washed in PBS and mounted with a mounting solution containing DAPI prior to observation using a fluorescent microscope. For counting the Inclusion bodies containing IBAGs, and the IBAGs within inclusion bodies in A3.01 and HEp-2 cells, the immunofluorescence images from three independent experiments were subjected to a manual counting. It was considered IBAGs all the clear distinct granular structures within the inclusion bodies.

### Real-Time PCR

HEp-2 and A3.01 cells inoculated with HRSV for 1 h at 4°C, were centrifuged at 200 × g, and part of the cells and supernatant were placed into Trizol reagent, which was considered as the time point zero. The remaining cells were incubated in CO_2_ at 37°C, and aliquots of cells and supernatant were collected in Trizol reagent at 1, 3, 6, 12 and 24 hours post-infection. The total RNA was isolated following the Trizol protocol. Afterward, cDNAs from total RNAs were obtained by reverse transcription with Superscript III Reverse Transcriptase (Invitrogen) primed with random primers. The real-time PCR assays were performed using primers targeting the gene for the N protein of HRSV. RNA quantification was extrapolated from a standard curve made with dilutions of plasmid containing the cloned RSV N gene segment.

### R18 Fluorescence conjugation and dequenching assay

These protocols were the same used by Evelyn M. Covés-Datson et al [32] with slight modifications. HRSV- or culture supernants of HEp-2 cells were incubated with 30 µg/mL of octadecyl rhodamine B chloride (R18) for 1 hour at room temperature protected from light. Then, fluorescently labeled R18-HRSV or culture supernatants of HEp-2 were separated from excess R18 by a separation column (PD-10 Desalting Columns GE Healthcare). HRSV-R18 or culture supernatants of HEp-2 were incubated in suspension with A3.01 and HEp-2 cells at 4°C for 1 h to allow virus attachment, and then incubated in a CO_2_ at 37°C, and the dequenching of R18 was measured in a Synergy Photometer at the appropriate time points. As positive control, 1% triton X-100 in PBS was used for total dequenching. The amount of fluorescence emitted from R18 was measured at 560 nm excitation and 590 nm emission.

### Fluorescent Focus Assay

A3.01 cells were infected with HRSV, and at 1, 6, 12, 24 and 48 hours post-infection, their supernatants were collected to investigate the progeny production of the HRSV in these cells. Serial 10-fold dilutions of the supernatants collected at each time point was made. Then, a HEp-2 cell monolayer was infected with each of the HRSV dilutions. Three days after the infection, the cells were permeabilized with 0,01% of 100X Triton and incubated with an antibody made in mouse anti-HRSV F protein conjugated with Alexa Fluor 488. The fluorescent focus were analyzed in a fluorescent microscope; the number of focus were counted and plotted in a graph.

## Results

### A3.01 lymphocytes inoculated with HRSV are inefficient in progeny production

The A3.01 cells were infected in suspension with HRSV (MOI=1) and analyzed by flow-cytometry at several times after infection (figure 1A). Infection was reproducibly detected in three independent experiments at all times post infection, with peak at 48 hpi, when over 40% of the cells were infected. The highest numbers of cells positive for HRSV N protein were found at 48 hpi (figure 1A). These results indicate that at least under the condition used in these experiments, A3.01 cells are susceptible to HRSV infection and produce viral protein N. A3.01 cells were permissive for HRSV replication; however, virus replication in this cell type was markedly reduced and delayed in comparison with HEp-2 cells, as indicated by quantification of the HRSV genome released into the culture supernatants (figure 1B). The fluorescent focus assay with mouse anti-HRSV F antibody indicated that infectious HRSV progeny production in A3.01 is also inefficient, with a replicative burst of less than one log_10_ from 6 to 48 hours post-infection (figure 1C).

**1.**
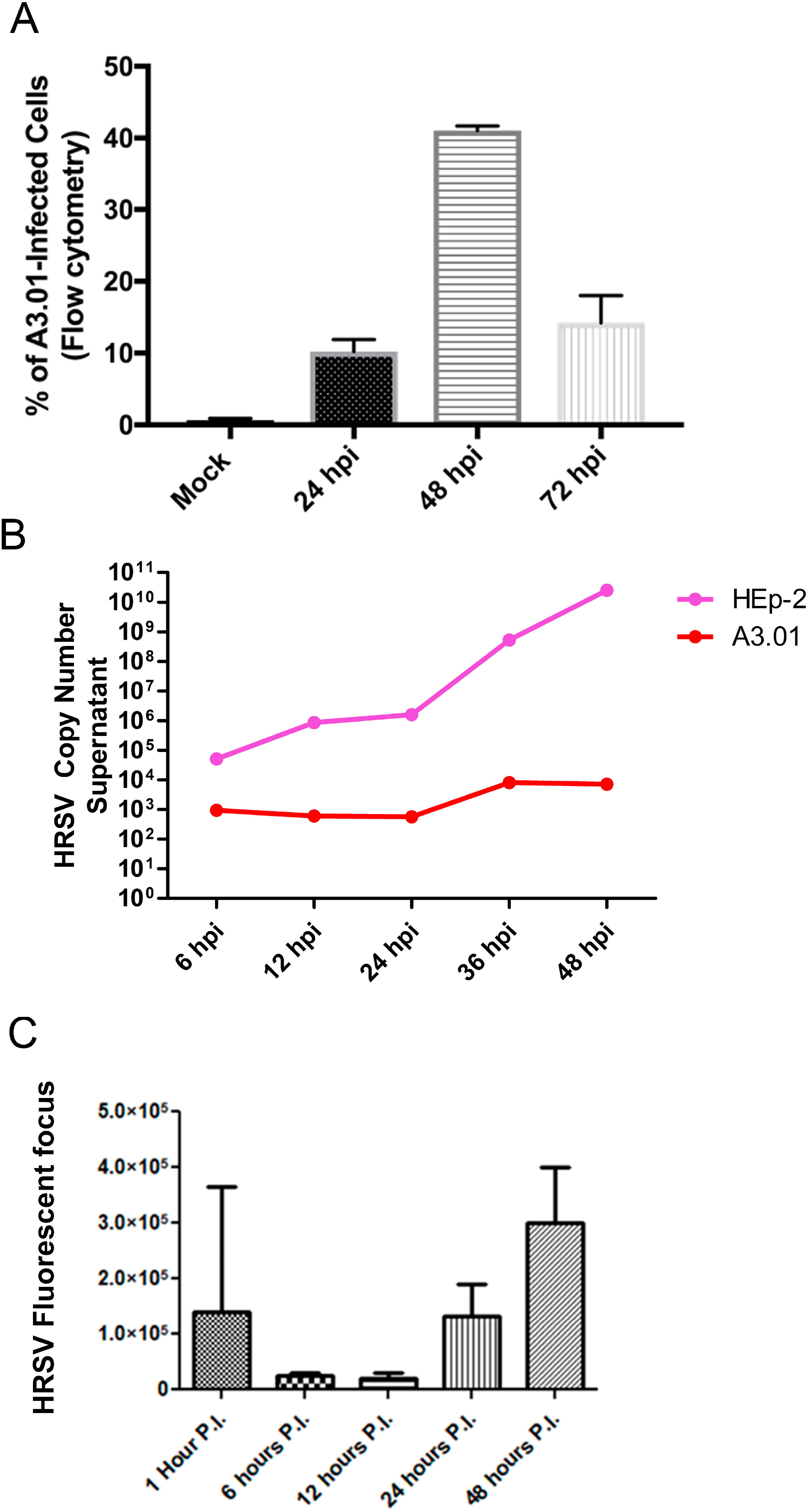
Infection of HRSV in A3.01 cells. (A) flow cytometry analysis of mock and HRSV-infected cells showing the percentage of the infected cells by detection of HRSV N protein. (B) qPCR of HRSV genome in supernatant of infected HEp-2 and A3.01 cells over time post-infection. (C) HRSV progeny production in A3.01 cells determined by fluorescent focus assay. All results are from three independent experiments.

### HRSV genome replication in A3.01 cells is inefficient

Since we observed that A3.01 cells were inefficient in progeny production of HRSV, we sought to investigate which step of the viral replicative cycle was compromised in these cells. We set out to assess the virus genome production by quantitative real-time PCR targeting the HRSV N gene in HRSV-infected A3.01 and HEp-2 cells, the latter of which was cultured either as cells attached to plates (Att) on in suspension (Sus). Cells and viruses were incubated for absorption for 1 hour at 4° C, the inoculum was washed away, and the cells were further incubated at 37°C. At several times post infection, cells were collected and processed for RNA extraction in Trizol. The same amount of virus inoculum was placed in virus-only wells, without cells, for quantification of the remaining virus inoculum after the incubation periods. The results revealed that the attachment of HRSV to HEp-2 and A3.01 cells was only slightly different, but the virus replication was almost 4 log_10_ higher in HEp-2 than in A3.01 cells (figure 2). Furthermore, the amount of cell-associated HRSV RNA decreased in A3.01 cells over 3 hours after inoculation, suggesting a failure in a step after virus-cell attachment such as virus-cell fusion, which would result in degradation of internalized virions.

**2.**
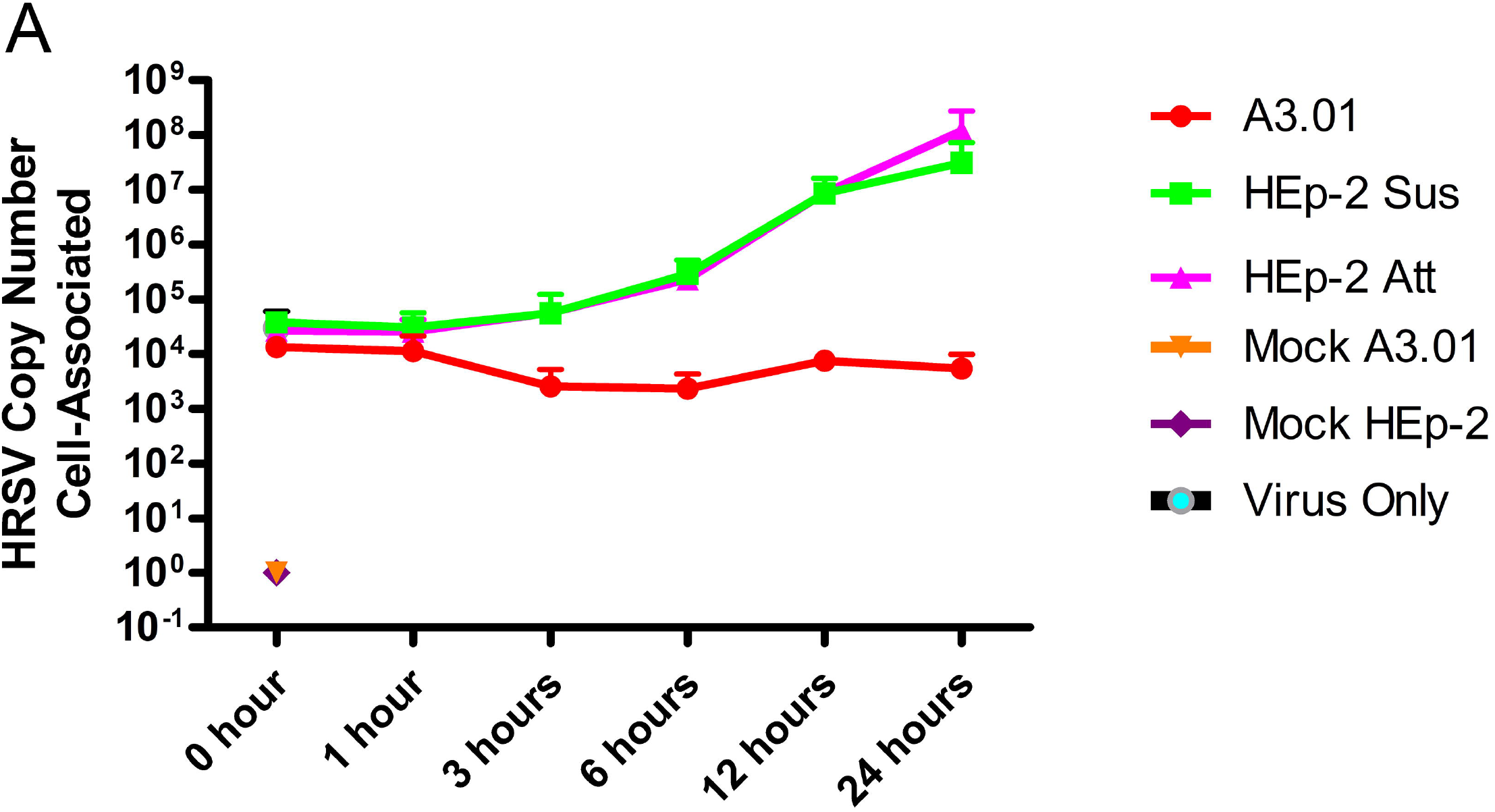
Intracellular accumulation of HRSV genome in A3.01 and HEp-2 cells. (A) A3.01 and HEp-2 cells attached (Att) or in suspension (Sus) were inoculated with HRSV or mock-inoculated, and kept at 4oC for 1 hour for attachment, then cells were centrifuged, and collected for qPCR analysis at time zero and at different times thereafter. Genome quantification was plotted in Y axis. The “virus only” well received only virus, in the absence of cells. This graph is a representation of three independent experiments.

### The fusion process of HRSV in A3.01 cells is defective

The numbers of HRSV-positive A3.01 cells by flow cytometry were dependent on the duration of the incubation period for virus adsorption to the cells (figure 3A). The fraction of cells positive for HRSV N protein at 24 and 48 hpi was higher when the virus inoculum was not removed than when it was washed away after 3 hours. This is consistent with the possibility that the entry of HRSV in A3.01 cells is not efficient. To test this possibility, both A3.01 and HEp-2 cell suspensions were inoculated with R18-labelled HRSV (R18-HRSV) or mixed with culture supernatants of mock-infected cells containing an equivalent amount of R18, and the virus-cell fusion was evaluated based on dequenching of R18, which occurs when viral envelope fuses with cellular membranes. We examined virus-cell fusion with 4×10^3^ cells per well. The virus or mock inocula were incubated with cells on ice for 1 hour to allow adsorption but not entry, after which the unbounded viruses were washed away. Subsequently, the virus-cell suspensions were placed at 37°C, and the fluorescent emission by the R18 de-quenching process was measured. The results revealed that at 1, 3, 6, and 24 hpi, the quantity of de-quenched R18 was significantly higher in HEp-2 than in A3.01 cells (figure 3B). To test the possibility that the difference in the de-quenching of R18 between the two cell types is due to the difference in the amounts of attached virus, we compared the attachment of R18-HRSV to HEp-2 and A3.01 cells using the same approach. Both A3.01 and HEp-2 cells were treated with 1% Triton X-100 immediately after the washes with cold PBS, thereby completely de-quenching the attached viruses (figure 3C). The results suggested that the amounts of HRSV attached in the A3.01 and HEp-2 cells are similar (figure 3C). Altogether, we conclude that the process of HRSV fusion to A3.01 cells is significantly compromised in comparison to HEp-2 cells.

**3.**
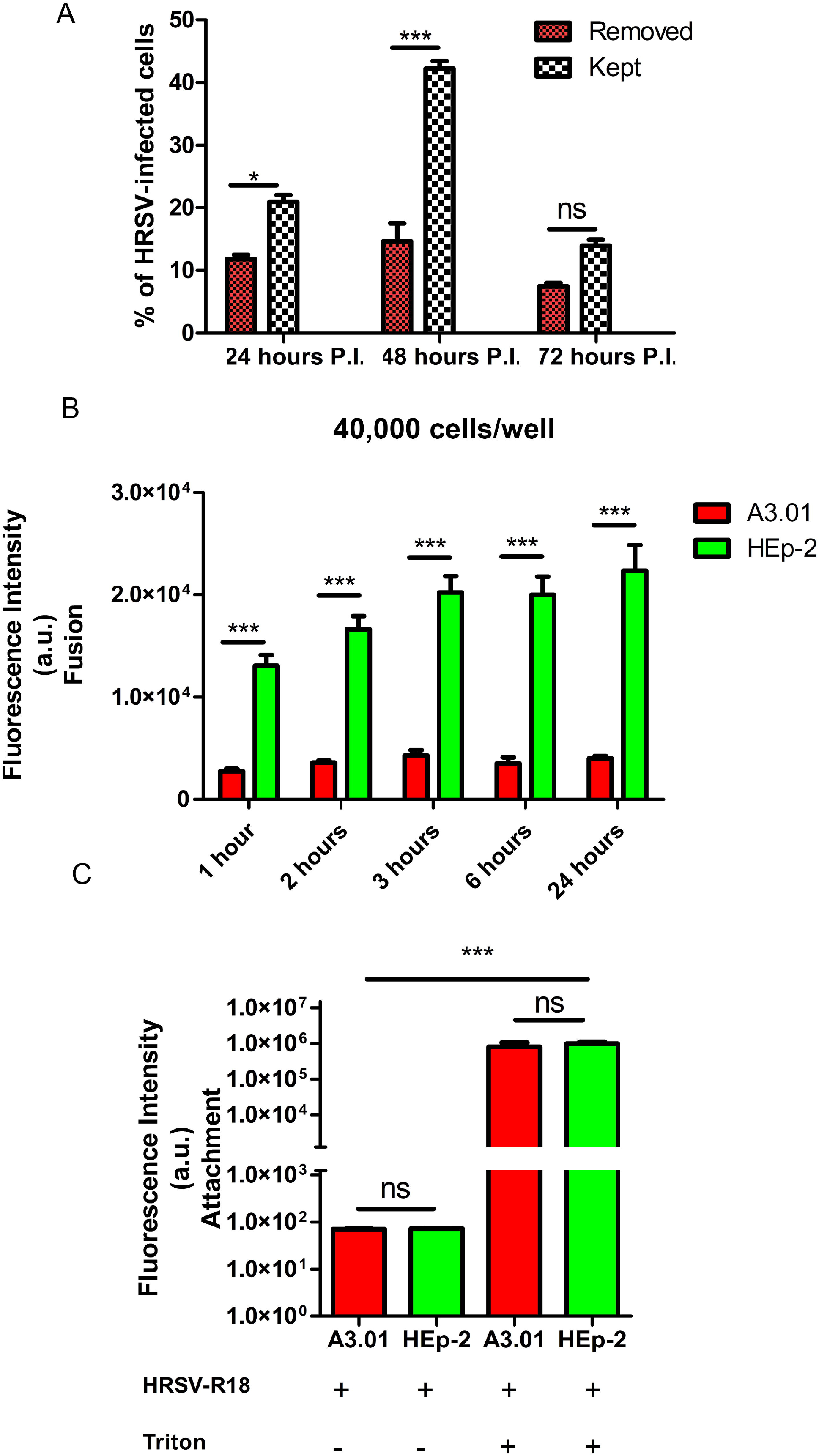
The fusion process in A3.01 is inefficient. (A) differences between HRSV-infected cells with and without inoculum removal at different times post-infection. (B) comparison of the fusion process in A3.01 and HEp-2 cells over time, in less populated, 40,000 cells per well. (C) comparison of the HRSV attachment in A3.01 and HEp-2 cells. The graphs in A, B and C represent at least three independent experiments; the statistical method used was Student′s T-test, p*<0.05, p**< 0.01, and p***< 0.001. The intensity of fluorescence emitted by R18 was measured by a SynergyTM Multi-Mode Microplate Reader.

### HRSV protein production in A3.01 cells is small

Even though the inefficient fusion process of the HRSV in A3.01 cells by itself could explain the dramatic differences between genome replication in A3.01 and HEp-2, we also hypothesized that other steps of the HRSV production could also be affected. Since the replication of HRSV in A3.01 cells was inefficient, we examined whether the production of N protein, a main component of virus factories, was also compromised in A3.01 cells. Mean fluorescence intensity analysis of infected HEp-2 and A3.01 cells at different time points using flow cytometry (figure 4A), showed that the N protein level in each HRSV-infected A3.01 cell was significantly lower than that in each HEp-2-infected cell at all times. With the greatest difference at 48 hpi when the quantity of protein in HEp-2 cells was approximately 42 times that found in A3.01 cells (figure 4B). Western blot analysis of HRSV proteins produced in A3.01 or HEp-2 cells revealed that not only the N protein but also the other viral proteins were less abundant in infected A3.01 cells at different times post-infection (figure 4C). Although the western blotting analysis is not controlled for the number of infected cells, it revealed that the fold differences in M and P levels between HEp-2 and A3.01 are even greater than the difference in N. Therefore, not only N, which was examined in the flow cytometry, but also other viral proteins are present in A3.01 at lower levels than in HEp-2. In addition, it was not even possible to see the bands corresponding to the HRSV G and F glycoproteins. These data indicate that the HRSV protein levels at different times post-infection in A3.01 are broadly compromised.

**4.**
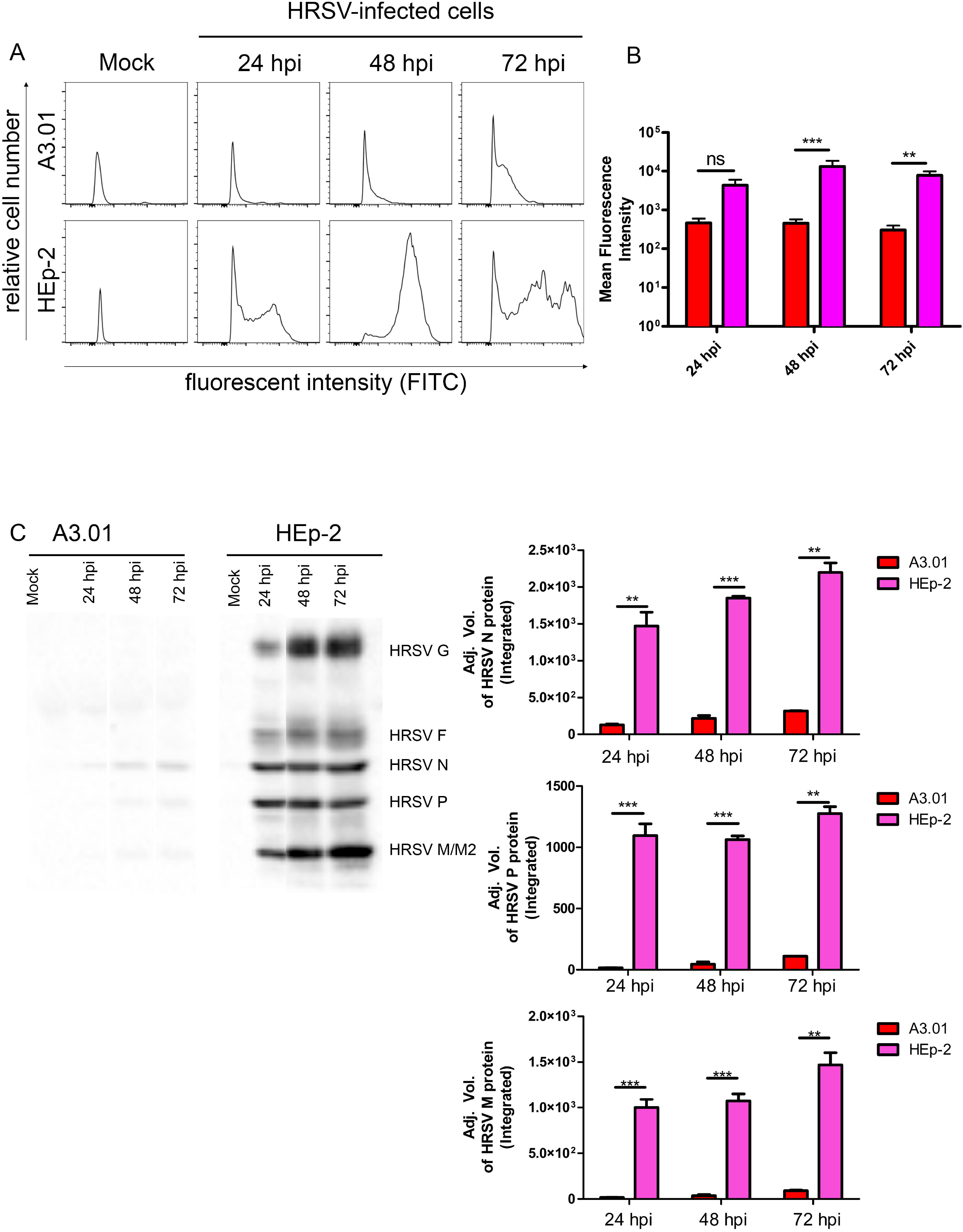
HRSV protein production in A3.01 cells is discrete. (A) histogram of mean intensities of fluorescence of cells by a flow cytometry. (B) graph plotted from three different experiments comparing the production of HRSV N protein in A3.01 and HEp-2 attached (Att) cells at 24, 48 and 72 hours post-infection. The graph in B represents at least three independent experiments; (C) western blot of HEp-2 and A3.01 cells infected or not (MOCK) by HRSV, the graphs represent the analysis of the proteins bands of three independent experiments. The statistical method used in figure B was Two-Way ANOVA, and the statistical method used in C was Student′s T-test, p*<0.05, p**< 0.01, and p***< 0.001.

### HRSV inclusion body formation is compromised in A3.01

The inclusion bodies (IB’s) are important platforms for HRSV replication and assembly [5,16,17]. Considering that the HRSV N, P and M are pivotal components of the IB formation and that the levels of these proteins are very low in A3.01 cells, we expected that the capacity of the HRSV to produce IBs in A3.01 cells is highly diminished. Interestingly, immunofluorescence for HRSV N protein revealed that HRSV infection does induce inclusion body formation in A3.01 cells (figures 5B and C). However, by measuring the perimeters of IBs in A3.01, we found that they were smaller than those in Hep-2 cells (figure 5G) and also present in significantly lower quantities in A3.01 compared to HEp-2 cells (figure 5H). Since the area of the HEp-2 cell is approximately 3.2 times larger than A3.01 cells, the difference in the area sizes can partially account for the differences found in the quantity of HRSV N inclusion bodies in HEp-2 and A3.01. However, even when we normalized the quantity of HRSV N inclusions by the area, the quantity of HRSV N-positive IBs was higher in HEp-2 than A3.01 cells (figure 5I).

**5.**
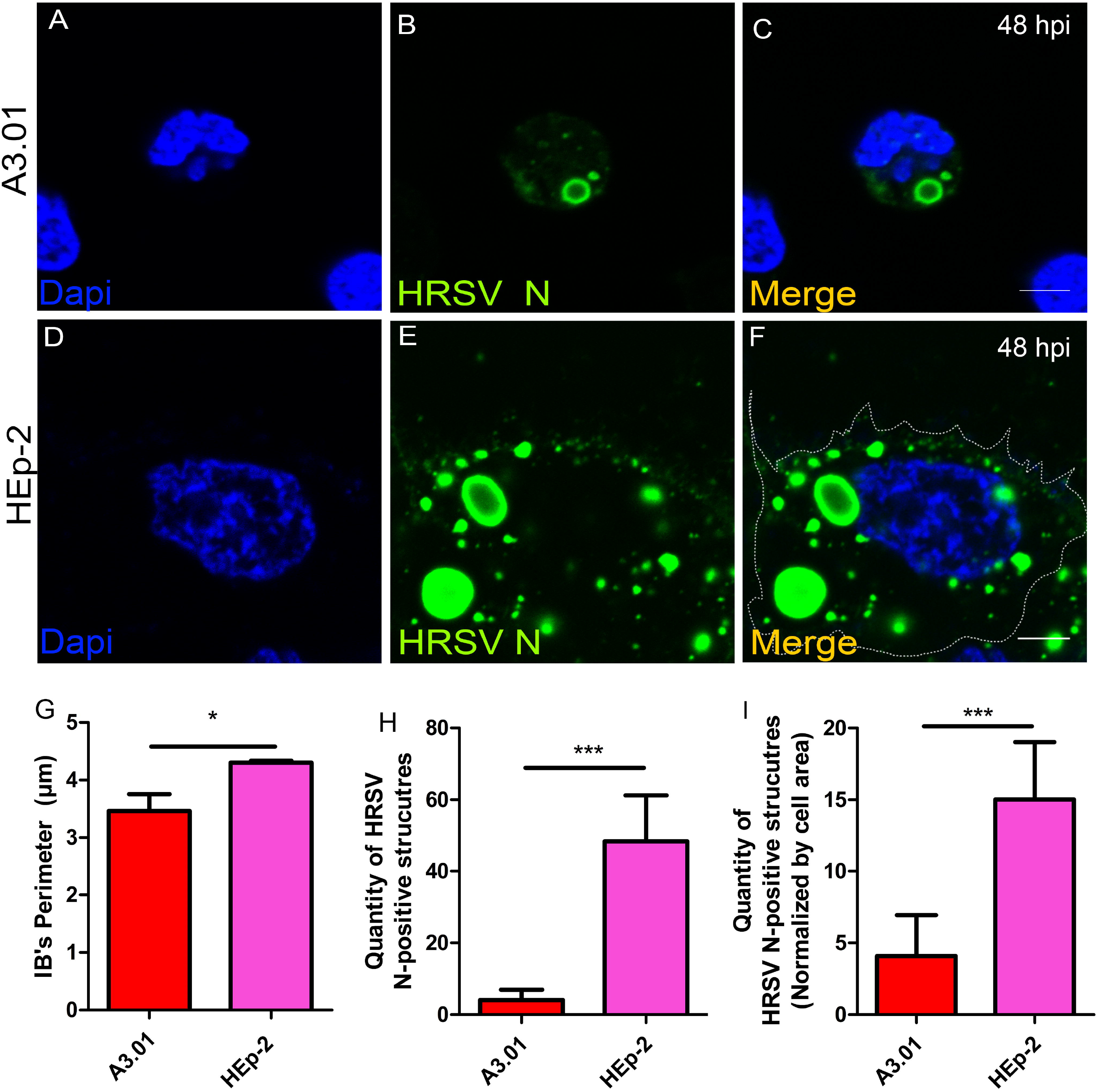
HRSV inclusion bodies in HRSV-infected A3.01 and HEp-2 cells. (A-C) A3.01 cells stained for HRSV N protein at 48 hpi (green fluorescence). (D-F) staining for HRSV N protein in HEp-2 cells at 48hpi. (G) comparative analysis of inclusion body sizes in A3.01 and HEp-2 cells. (H) comparative analysis of numbers of vesicular structures stained for HRSV N in A3.01 and HEp-2 cells. (I) comparative analysis of the structures stained for HRSV N in A3.01 and HEp-2 cells normalized by cell area. The immunofluorescence images shown in figures A-F represent a single focal plane of at least three independent experiments. The images were taken in a Zeiss 780 confocal microscope. Magnification 63x. The scale bars represent = 10µm. The graphs in G and H represent at least three independent experiments; the statistical method used was Student′s T-test, p*<0.05, p**< 0.01, and p***< 0.001.

### The inclusion bodies of HRSV in A3.01 cells lack IBAGs

Recently, Rincheval et al [14] showed that HRSV inclusion bodies are compartmentalized and contain ultrastructural granules, which were called inclusion body-associated granules (IBAGs). They also proposed that these structures are important for the IB’s functionality, since this organization promotes stabilization of viral transcripts and hence an efficient protein production process. We performed a FISH analysis to examine the formation of IBAGs of HRSV-infected A3.01 cells. A biotinylated poly-T probe was used to reveal regions enriched for polyadenylated RNA. HRSV produced IBAGs at both 24 and 48 hpi in HEp-2 cells (Figures 6I-L, and M-P), but in A3.01 cells the majority of the HRSV-induced IBs did not appear to contain prominent IBAG’s and therefore can be considered hypofunctional (figures 6 Q-T and U-Z pointed by arrowheads). HRSV IBs in A3.01 cells did contain staining for polyA, but such polyA signal was rarely found as clear distinct granules within the inclusion bodies as observed in HEp-2 cells. A Z-stack and quantitative analyze was performed between labelled HEp-2 (supplementary figure 1A) and A3.01 cells (supplementary figure 1B). The quantity of inclusion bodies of A3.01 and HEp-2 cells containing distinct IBAG’s within were counted (supplementary figure 1C). It showed that the quantity of inclusion bodies of infected A3.01 cells containing IBAG’s is significantly lower than those in infected HEp-2 cells (supplementary figure 1C). Further, the quantity of the IBAGs found in inclusion bodies was significantly lower in A3.01 cells than in HEp-2 cells (supplementary figure 1D). Therefore, we conclude that even when poly A-containing mRNAs are present in the IBs of HRSV-infected A3.01 cells, the sequestration of such mRNAs to subcompartments of IBs is inefficient in this cell type.

**6.**
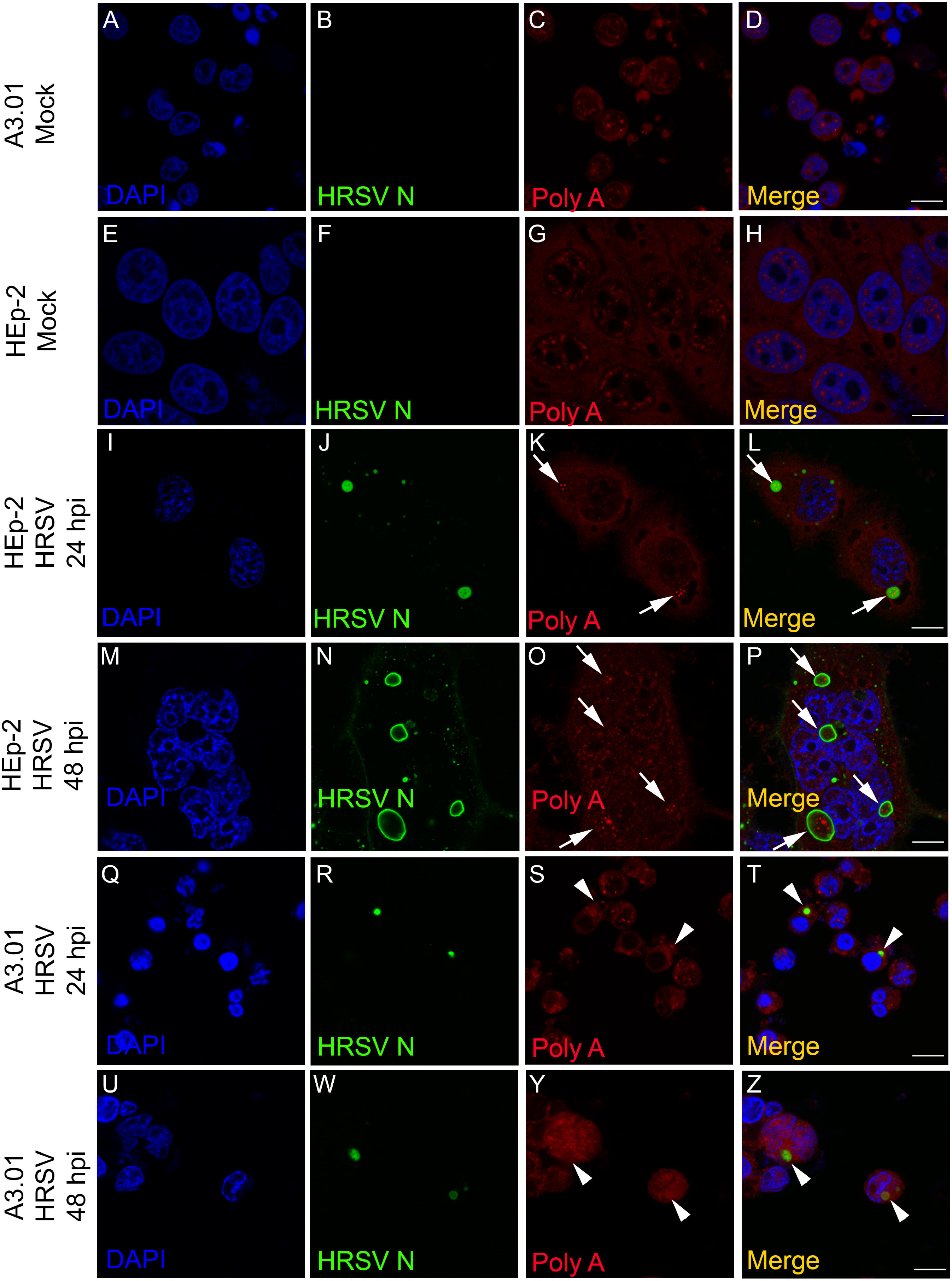
The HRSV inclusion bodies in A3.01 are IBAG’s absent. (A-D) and (E-H) A3.01 and HEp-2 mock-infected cells. (I-L) and (M-P) HRSV infected HEp-2 cells at 24 and 48 hpi as shown by IBs in J and N, respectively. Within IBs in J and N it is possible to see FISH-signal for IBAGs, as shown in K, L, O and P (arrows). (Q-T) and (U-Z) represent A3.01-infected cells at 24 and 48 hpi. No IBAGs were seen within HRSV inclusion bodies (R and W) in these cells, as shown in S, T, Y and Z. This set of figures represents a single focal plane of three independent experiments taken in a Zeiss 780 Confocal. Magnification 63x. Scale bars = 10µm.

### The HRSV F protein partially co-localizes with Golgi markers giantin and TGN46 in A3.01 cells

In epithelial cells, F protein, a transmembrane protein, follows the conventional anterograde pathway from ER through Golgi to the plasma membrane [36]. Also, the Golgi participates in the intracellular traffic of the HRSV N protein to the plasma membrane [23,24]. To examine whether the trafficking of HRSV F and N proteins in A3.01 lymphocytes is similar to the one in HEp-2 cells, immunofluorescence staining was done for a cis- and medial-Golgi marker, giantin, a trans-Golgi network marker, TGN46, and the viral proteins F and N. A partial co-localization of HRSV F protein with the Golgi markers was observed at 48 hours post-infection (Figure 7E-H). In HEp-2 cells, the HRSV F protein shows strong colocalization with Giantin and partial co-localization with TGN46 (supplementary figures 2A-F). Even though the HRSV N protein is a cytoplasmic and not transmembrane protein, a partial colocalization with TGN46 was also observed in HEp-2 cells (supplementary figures 3A-F). However, in A3.01 cells, the N protein did not co-localize with the Golgi markers (Figure 7 I-L). Therefore, we conclude that the HRSV F and N proteins colocalized with the Golgi in A3.01 cells to a lesser extent than in HEp-2 cells.

**7.**
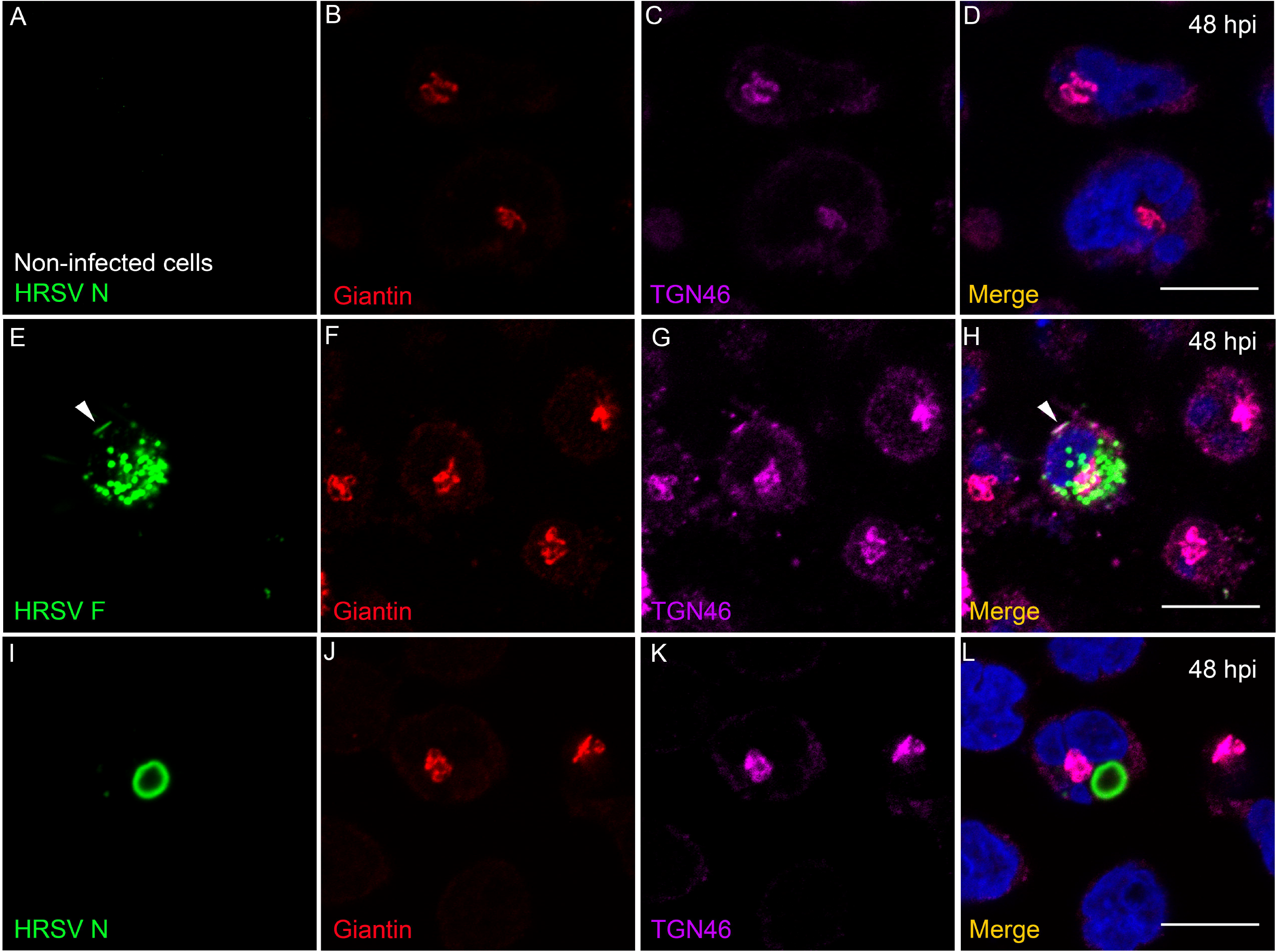
Co-localization analysis of HRSV proteins at the Golgi in A3.01 cells. (A-D) mock-infected cells stained for cis and medial-Golgi (Giantin) in red (B) trans-Golgi (TGN46) in magenta (C) merge in (D). (E-H) HRSV-infected A3.01 cells at 48 hpi stained for HRSV F (green) in €, giantin in (F) and TGN46 in (G). Merge is depicted in (H). The arrowheads point to a region where the F protein is located in the plasma membrane. (I-L) HRSV-infected A3.01 cells at 48 hpi stained for HRSV N in (I), giantin in (J) and TGN46 in (K). The merge of the figure is in (L). All the figures represent a single focal plane of at least three independent experiments taken in a Zeiss 780 Confocal microscope. Magnification 63x. Scale bars = 10µm.

### The HRSV F and N proteins partially co-localize with markers of endosomal pathway

After trafficking through the Golgi stacks, the transmembrane proteins often associate with endosomal pathways [33–35]. In addition, it is already known that some of the HRSV proteins follow the endosomal system to reach the plasma membrane [19,23,24,36]. To investigate whether the HRSV proteins also associate with endosomal machineries in A3.01 cells, immunofluorescence assays were performed for several markers of this pathway. First, we aimed to investigate whether HRSV proteins were preferentially targeted to late/lysosomal pathway in these cells. We assessed HRSV proteins co-localization with early endosome marker, the Early Endosome Antigen 1 (EEA1), and with a lysosome-specific marker, LAMP-1 (figure 8). Indeed, there was strong co-localization of Lamp-1 signal with HRSV proteins in A3.01 cells (figures 8E-L) when compared to HEp-2 (supplementary figures 2 and 3M-O, pointed by arrows), suggesting an involvement of lysosomes in protein trafficking or degradation. However, the imaging evidence is not enough to confirm specific involvement of lysosomal machinery in HRSV protein degradation, since the co-localizations of HRSV proteins with Lamp-1 and EEA1, an early endosome marker, were not statistically different in A3.01 cells (figures 8M and N). We also did immunofluorescence microscopy for CD63 and Sorting Nexin 2 (SNX2). CD63 is a marker for late endosomes, while SNX2 is part of the cellular retromer complex, responsible for carrying cargoes from early endosomes to the trans-Golgi [37,38]. In addition, recently, it was found that SNX2 is recruited to HRSV N inclusion bodies and plays a role in the HRSV viral production in HEp-2 cells [24]. It was possible to see that, similarly to what happens in HEp-2 cells, the HRSV F protein partially co-localized with SNX2 and CD63 in A3.01 (figures 9I to L, and supplementary figures 2G to L). In addition, the HRSV N protein was found juxtaposed to CD63 both in A3.01 and HEp-2 cells (figures 9I to L, and supplementary figures 3J to L, pointed by arrows). Interestingly, however, in contrast to HEp-2 cells, the HRSV N-positive structures failed to recruit SNX2 in A3.01 cells (figures 9E to H and supplementary figures 3G to I).

**8.**
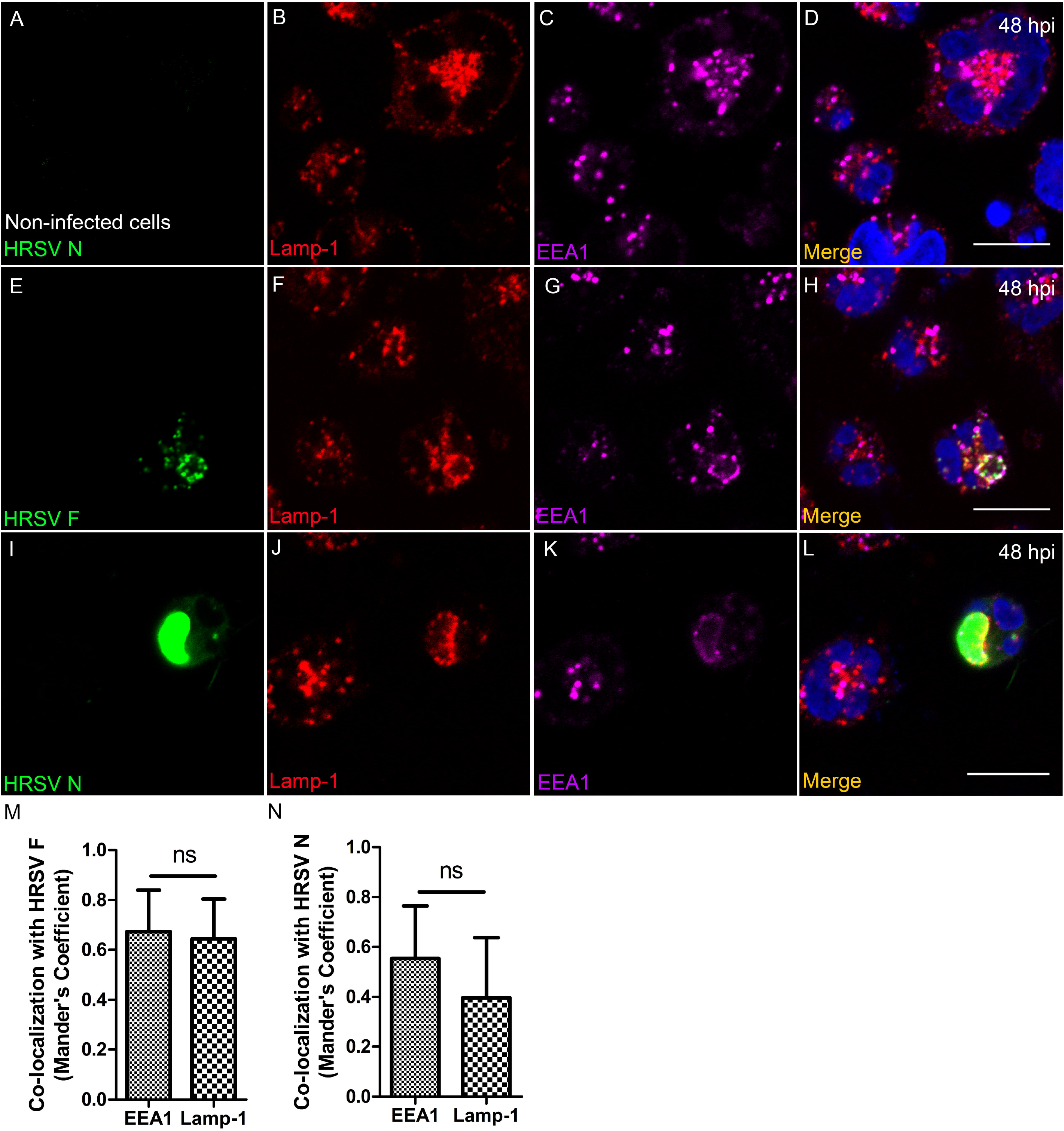
Co-localization of EEA1 and Lamp-1 with HRSV proteins in A3.01 cells. A-D, represent A3.01 mock-infected cells. In B is showed Lamp-1 (red), in C EEA1 (magenta) and in D the merge of the set of the figures. E-H, correspond to A3.01-infected cells, stained by HRSV F, shown in E, Lamp-1 in F and EEA1 in G. The merge to this set of figures is depicted in H. I-L, correspond to A3.01-infected cells, stained by HRSV N, shown in I, CD63 in J and SNX2 in K. The merge of this figure set is depicted in L. All the images were taken in a Zeiss 780 Confocal and are a representation of a single focal plane. Magnification 63x. Scale bars = 10µm.

**9.**
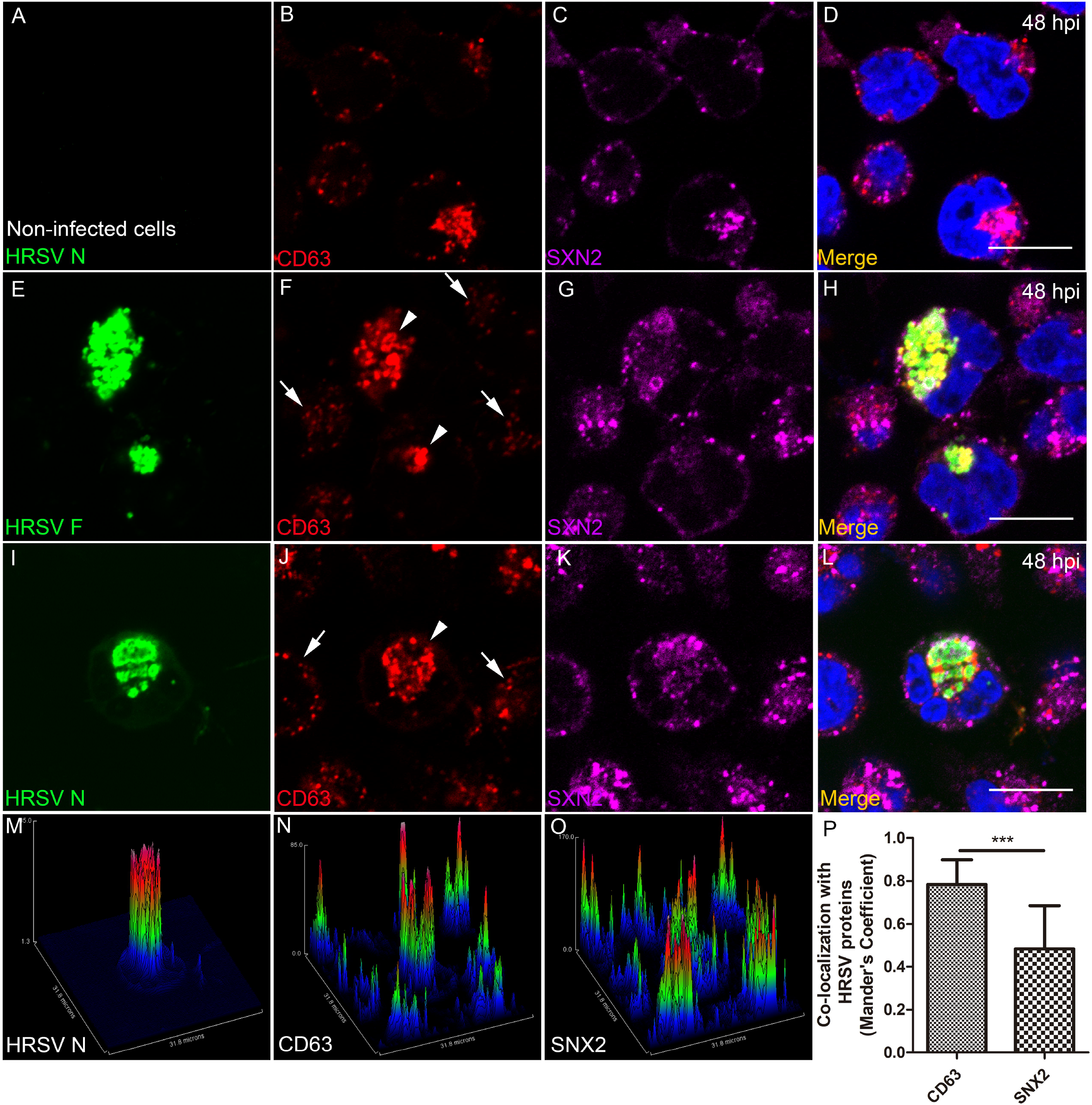
CD63 co-localization with HRSV proteins. (A-D), represent A3.01 mock-infected cells. In B is showed CD63 (red), in C SNX2 (magenta) and in D the merge. E-H, correspond to A3.01-infected cells, stained by HRSV F, shown in E, CD63 in F and SNX2 in G. The merge to this set of figures is depicted in H. The arrows point to the places where the cells are not infected, and there were not CD63 accumulation. The arrowhead point to the places where the HRSV F protein is found and the CD63 is accumulated. I-L, correspond to A3.01-infected cells, stained by HRSV N, shown in I, CD63 in J and SNX2 in K. The merge to this set of figures is depicted in L. The arrows point to the places where the cells are not infected, and there were not CD63 accumulation. The arrowhead points to the place where the HRSV N protein is found, and the CD63 is accumulated. M-O represent the surface plot of the figures in I, J and K respectively showing the heating map of the CD63 where HRSV N is located. P, graph of co-localization between CD63 and SNX2 with HRSV proteins, showing significant co-localization of CD63. The figure P represents more than three independent experiments and was done in at least five cells per field, the statistical method used was Student′s T-test, p*<0.05, p**< 0.01, and p***< 0.001. All the images were taken in a Zeiss 780 Confocal and are a representation of a single focal plane. Magnification 63x. Scale bars = 10µm.

### The production of HRSV filaments at the plasma membrane of A3.01 cells is very low

HRSV assembly/budding takes place at the plasma membrane, with the appearance of viral protein-containing filament-shaped structures in infected epithelial cells [36,39].

Since the filaments emerging from the plasma membrane is one of the hallmarks of the HRSV infection [1], we examined whether HRSV is capable of producing typical filamentous structures in A3.01 cells. Immunofluorescence for viral proteins was done with special attention to the plasma membrane. Although only scarcely, HRSV N and M proteins could be seen in filament-shaped structures (figures 10A-C, arrowheads). Nevertheless, the quantity of filament structures emerging from A3.01 cells is minimal when compared to those found in HEp-2 cells (figures 10D-J). It is noteworthy that not only the filament formation is reduced in A3.01 cells (figures 10A-C and H-I), the quantity of the A3.01 cells displaying at least one filament is significantly lower than that of HEp-2-infected cells (figure 10G). Therefore, while HRSV proteins were readily detected in intracellular compartments of A3.01 cells, very little accumulation of HRSV products was seen at the plasma membrane, which results in a fewer filament production in these cells.

**10.**
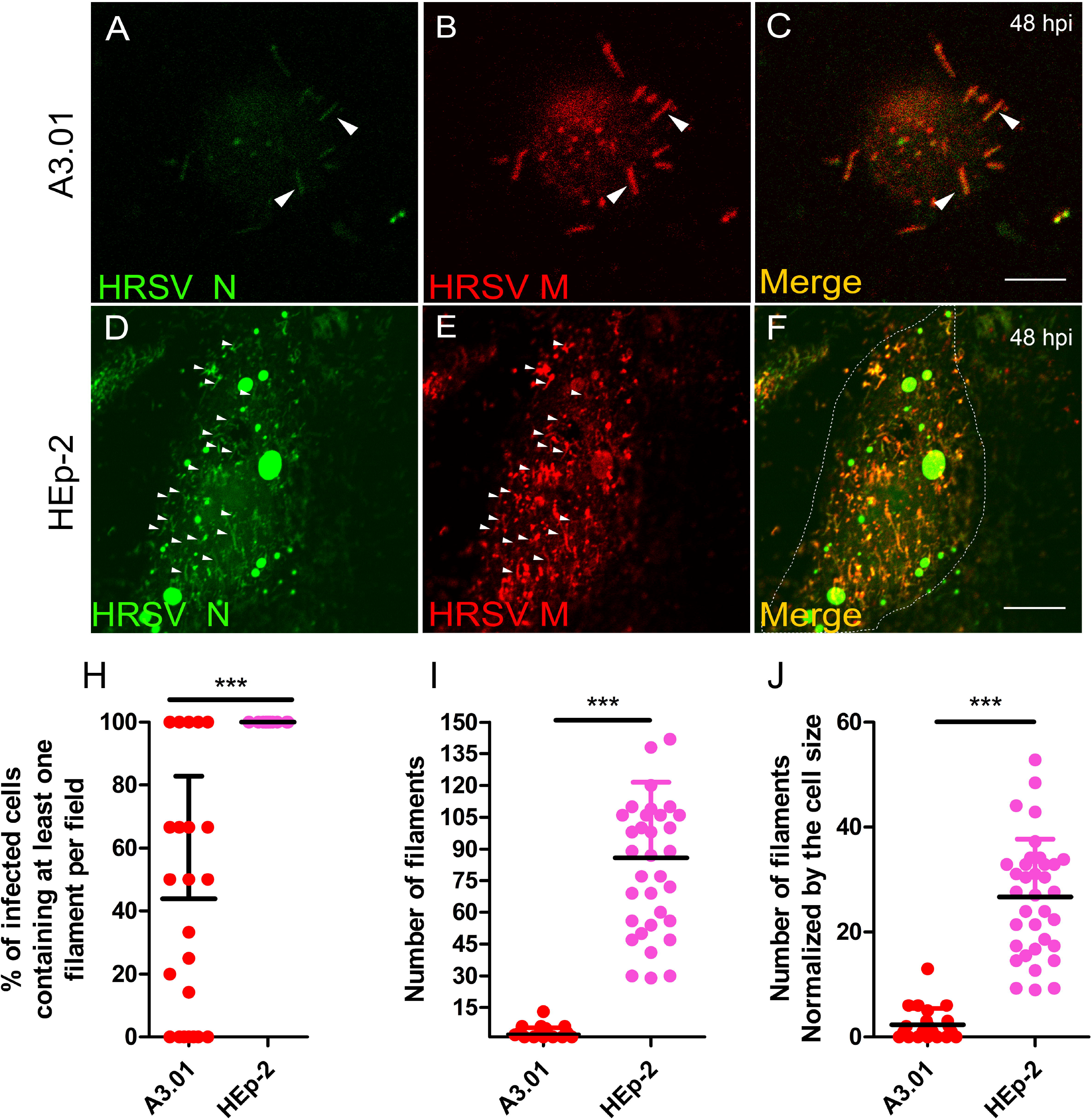
Filament formation of HRSV in A3.01 is rare. (A-C) immunofluorescence for HRSV N and M in A3.01-infected cells at 48 hpi depicting some filaments, pointed by arrowheads. (D-F) immunofluorescence for HRSV N and M in HEp-2-infected cells at 48 hpi depicting filaments pointed by arrowheads. Figures A to F represent a single focal plane of at least three independent experiments taken in a Leica SP5 Confocal. Magnification 63x. This experiment was repeated at least three independent times. The scale bar of figure C = 10µm. (H) graph of the percentage of A3.01 and HEp-2-infected cells displaying at least one filament emerging from the plasma membrane per field. (I) graph of the quantity of the filaments in A3.01 and HEp-2-infected cells. (J) graph of the quantity of the filaments in A3.01 and HEp-2-infected cells normalized by the cell area. The graphs depicted in H, I and J are representative of more than 5 independent experiments. The statistical method used was Student′s T-test, p*<0.05, p**< 0.01, and p***< 0.001. All the images were taken in a Zeiss 780 or Leica SP5 Confocal and are a representation of a single focal plane. Magnification 63x. Scale bars = 10µm.

## Discussion

Studies on the interaction of HRSV with host cells are usually conducted using respiratory epithelial cells, which are the main targets of the natural infection that usually results in cell death [1]. However, HRSV is also able to infect non-respiratory cells, including CD4^+^ T lymphocytes [2,3,5], and may cause persistence in some cell types, like murine macrophages [40]. The frequent detection of HRSV RNA in tonsillar tissues from children without symptoms of acute HRSV infection [4] suggests that the agent may cause prolonged infection in secondary lymphoid tissues. It is thus presumable that HRSV causes patterns of infection that differ between lymphoid and epithelial cells, which are likely to affect functions and survival of these cells. The present study was undertaken to elucidate details of in vitro HRSV infection of the human CD4^+^ T cell line A3.01 in comparison to infection of HRSV in commonly used cell line, HEp-2. We showed that HRSV infection in A3.01 cells is different from that in HEp-2 cells at multiple steps of the virus replication cycle.

While HRSV was able to infect A3.01 cells, the progeny production in these cells was much lower than that of Hep-2 cells. This could be due to reduced susceptibility and/or permissiveness of A3.01 cells to infection by HRSV. In that regard, we showed that HRSV fusion with A3.01 cells was reduced as compared to the HEp-2 cells. This could be due to some limitation of the HRSV fusion process itself or due to a difference in HRSV-receptor engagement in A3.01 cells, creating subsequent hindrance to the fusion process. It is noteworthy that HRSV can use different receptors to fuse to host cell membranes [10,41,42]. Therefore, while binding of HRSV to HEp-2 and A3.01 cells were comparable, differences in the levels of expression or affinity of individual receptors between the two cell types may affect the efficiency of virus-cell fusion. It is also worthwhile testing the activity of macropinocytosis in A3.01 cells versus HEp-2 cells, since this is the best known mechanism of HRSV internalization [9,43].

The replication of the HRSV genome occurs in IBs [1]. Recently Rincheval et al. demonstrated that the HRSV mRNAs are sequestered within the IBs, more specifically in organized structures that they called inclusion bodies-associated granules or IBAGs [14]. Using IF and FISH approaches in the present study, we found that HRSV IBs in A3.01 cells are significantly less abundant, smaller, and morphologically different than those seen in Hep-2 cells and that most of the IBs that are formed in A3.01 cells lack IBAGs. It is thought that in IBAGs, the M2-1 protein of the HRSV binds to viral mRNAs and thereby makes them more stable, which consequently ensures a better protein production [14]. Therefore, it is likely that the lack of discrete HRSV IBAGs in A3.01 cells at least partially accounts for the reduction in HRSV proteins N, M and P in A3.01 cells in comparison with HEp-2 cells at all times post infection (figure 4). HRSV proteins N and P are integral parts of the RNA transcription complex and IBs [44–46]. Therefore, lower levels of these proteins could help to explain significantly reduced rates of HRSV genome production in A3.01 cells compared to the HEp-2 cells. Together, these findings lead us to speculate that the low permissiveness to the HRSV genome replication in A3.01 cells is a sum of defects in formation of IBs and IBAGs and protein translation, where inefficiencies in each one of these steps exacerbate the other.

Since HRSV uses the secretory pathway to deliver viral proteins to the assembly sites at the plasma membrane [1,15,17,19,23,24], we examined the presence of virus proteins along the main components of the secretory and endosomal pathways by immunofluorescence. Our results and previous literature have shown that HRSV F and N proteins partially co-localized with markers of the secretory pathway [1,23,24]. However, it was noteworthy that in contrast to HEp-2 cells, A3.01 cells did not display an evident accumulation of SNX2 at places where the HRSV N protein is. In HEp-2 cells, SNX2 was found recruited to N-positive structures, and the knockdown of SNX2 impacted negatively in the HRSV production [24]. The absence of the recruitment of SNX2 to N-positive structures in A3.01 cells could contribute to the inefficiency in the viral production in A3.01 cells. The co-localization of CD63 and Lamp-1 with HRSV proteins is consistent with viral protein degradation, which could explain the lower amounts of the HRSV proteins in A3.01 cells. However, more studies should be performed to specifically assess this question, since it is currently unknown whether the viral protein synthesis is impaired or whether they are degraded in this cell type.

Interestingly, even though the HRSV proteins partially co-localized with secretory pathway markers in A3.01 cells, the number of the A3.01-infected cells displaying filaments at its surface was dramatically low. These results suggest that the trafficking of viral components to the virus assembly sites is defective in A3.01 relative to HEp-2 cells. We do not rule out at this time that the absence of viral filaments at the cell surface is due to a defect in the assembly process, which may be caused by low viral protein levels or by yet another block on the process at the plasma membrane.

Overall, the present results showed that HRSV infection of A3.01 CD4^+^ T cells is virtually unproductive, with the meager virion production as compared to HEp-2 cells, and that this is due to multiple defects during HRSV replication in A3.01 cells, namely, low virus-cell fusion, formation of hypo-functional inclusion bodies lacking IBAGs, failure to achieve high viral protein levels, and possibly altered trafficking of viral proteins and genome to virus assembly sites at the plasma membrane.

## Acknowledgements

We thank the expert technical support of Maria Dolores Seabra Ferreira and Jose Augusto Maulin (FMRP-USP Electron Microscopy Facility); Elizabete Rosa Milani and Roberta Ribeiro Costa Rosales (FMRP-USP Bioimaging Facility); and Maria Lucia Silva. This work was made possible by a research grant from the State of Sao Paulo Research Foundation (FAPESP, Grant No. 2013/16349-2). We also thank the funding agencies CNPq and CAPES (Brazilian Ministry of Education) for scholarships. We also would like to thank the Department of Microbiology and Immunology of the University of Michigan, and all the Ono’s Lab members for their precious suggestions. In addition, we thank all of the Eurico’s lab members for their pivotal ideas, experimental and discusses for the paper. Finally, we thank the Lukacss lab, especially Susan Morris for all the support and help.

## Legends for figures

**Supplementary Figure 1. Sectional slices from HEp-2 and A3.01-infected cells (supplementary)**. A, a Z stack was performed, and it is possible to observe IBAG’s within HRSV in HEp-2 IB’s along with the stacks, highlighted in the crops. B, sequential sectional slices from A3.01 cells, showing IB and the absence of clear IBAG’s within, highlighted in the crops. (C) graph of the quantity of the Inclusion Bodies containing IBAG’s found in A3.01 and HEp-2-infected cells at 48 hpi. The images were acquired in a Zeiss 780 Confocal, and are representative of three independent experiments. (D) graph of the quantity of IBAG’s counted in inclusion bodies in A3.01 and HEp-2-infected cells. Magnification 63x. Scale bars = 10µm. To the (C) and (D) 0graphs was counted at least 10 fields of three independent experiments from single focal plane of Z-stacks imaging acquisition. The statistical method used was Student′s T-test, p*<0.05, p**< 0.01, and p***< 0.001.

**Supplementary Figure 2. Colocalization of the HRSV Fusion protein with secretory pathway markers**. (A, B and C) immunofluorescence of HRSV F, Giantin and the merge respectively, the arrow points to colocalization. (D, E and F) immunofluorescence of HRSV F, TGN46 and the merge respectively, the arrow points to colocalization. (G, H and I) immunofluorescence of HRSV F, SNX2 and the merge respectively. (J, K and L) immunofluorescence of HRSV F, CD63 and the merge respectively. (M, N and O) immunofluorescence of HRSV F, Lamp1-1 and the merge respectively, the arrows point the areas of colocalization. All figures represent a single plane of Z-stack or not experiments. The images were acquired in a Zeiss 780 or Leica SP5Confocal. Magnification63x. Scale Bars = 10µm.

**Supplementary Figure 3. Colocalization of the HRSV Nucleoprotein protein with secretory pathway markers**. (A, B and C) immunofluorescence of HRSV N, Giantin and the merge respectively, the arrows point to colocalization area. (D, E and F) immunofluorescence of HRSV N, TGN46 and the merge respectively, the arrow points to colocalization. (G, H and I) immunofluorescence of HRSV N, SNX2 and the merge respectively, the arrows point to colocalization areas. (J, K and L) immunofluorescence of HRSV N, CD63 and the merge respectively, arrows point to colocalization areas. (M, N and O) immunofluorescence of HRSV N, Lamp1-1 and the merge respectively, the arrows point the areas of colocalization. All figures represent a single plane of Z-stack or not experiments. The images were acquired in a Zeiss 780 or Leica SP5Confocal. Magnification63x. Scale Bars = 10µm.

